# Cancer Diagnostics using Machine Learning of Tumor- and Tissue- Specific RNA Transcripts

**DOI:** 10.1101/2025.05.27.656256

**Authors:** Zhiao Chen, Jiawei Tang, Haochen Li, Yang Chen, Xianghuo He

## Abstract

Accurate cancer diagnosis and tissue origin identification are crucial for precision oncology. We explored the potential of tumor-specific RNA transcripts (Tumor-SRTs) and tissue-specific RNA transcripts (Tissue-SRTs) as dual biomarkers using machine learning. Tumor-SRTs effectively distinguished malignant from normal tissues across training, test, and validation sets. Classifiers trained on 25 Tissue-SRTs exhibited high performance, validated externally with varying predictive accuracy across tissue types. We developed SRT-based Cancer Diagnostics (SRT-CD), an intelligent diagnostic system, to diagnose primary and metastatic cancers and determine tissue origin of Cancers of Unknown Primary. SRT-CD achieved top-1/top-3 accuracies of 80%/98.1% for primary and 76.9%/92.3% for metastatic cancers in external validation. This study establishes SRT-CD as a robust tool for clinical cancer diagnosis and tissue origin identification, and guiding personalized treatment strategies.

## Introduction

RNA transcripts mediate genetic information flow from DNA to proteins through transcriptional and post-transcriptional regulation, generating extensive transcriptomic diversity. Over 95% of human genes produce multiple transcripts, contributing to substantial variation in gene expression ^1^. Different tissues preferentially express specific transcripts, leading to transcript-level differences, with selective transcription being a common feature across tissue types ^2,3^. Malignant tumors also exhibit diverse transcriptomes, each with unique transcript expression profiles ^4^. With the growing availability of RNA sequencing data from various tissues, cells, and tumors, large-scale comparative analyses of pan-tissue and pan-cancer datasets enable transcriptomic profiling and identification of tissue-specific RNA transcripts (Tissue-SRTs) or tumor-specific RNA transcripts (Tumor-SRTs) ^5-7^. In our previous work, we collected third-generation PacBio and ONT transcriptome sequencing data from 198 tumor samples and 153 normal samples to develop FLIBase, a comprehensive database of full-length annotated transcripts ^8^. This database covers multiple cancer types and normal tissues, identifying numerous previously unannotated Tumor-SRTs and providing detailed Tissue-SRT expression data. We define SRTs as transcripts uniquely expressed in particular tissues or diseases, including isoforms, variants, and novel transcripts. Tissue-SRTs are exclusive to normal tissues, while Tumor-SRTs are unique to tumors.

Tumor pathological diagnosis remains the gold standard for determining tumor nature, origin, staging, grading, and differentiation, offering critical insights into prognosis and treatment strategies. However, histologically similar tumors can complicate diagnosis, particularly in cases of Cancers of Unknown Primary (CUP), where the primary tumor site is unidentified ^9^. CUP exhibits aggressive clinical features, including early metastasis, rapid progression, and poor prognosis, with treatment largely dependent on conventional chemotherapy, which is often ineffective. Accurate identification of the primary tumor origin is crucial for targeted therapy decisions ^10^. Diagnosing CUP is particularly challenging due to nonspecific symptoms and multiple potential primary sites, necessitating a multidisciplinary approach for improved outcomes ^11^.

Machine learning, a subfield of artificial intelligence, applies statistical and computational techniques to analyze patterns within data, enabling accurate predictions and informed decision-making ^12^. In oncology, these techniques extract valuable insights from vast, complex datasets, thereby assisting healthcare professionals in achieving more precise and timely diagnoses, ultimately improving treatment outcomes and survival rates ^13-15^. In this study, we developed a machine learning-based cancer diagnostic system to classify cancer types and determine the tissue origins of CUP from RNA sequencing data. The approach involved identifying the most relevant features, followed by constructing and validating a diagnostic model based on the tumor- or tissue-SRTs. Our SRT-based Cancer Diagnostics (SRT-CD) system effectively classifies cancer types and identifies their tissue origins with high accuracy.

## Results

### Machine Learning of Tumor-SRTs for Precision Cancer Diagnosis

Our previous studies identified a subset of Tumor-SRTs exclusively expressed in tumor tissues ^8^. Here, we investigated whether these Tumor-SRTs could serve as reliable biomarkers for distinguishing tumor from normal tissues in matched human cancer samples. The first dataset was divided into a training and test set at a 3:1 ratio. Positive samples were derived from all The Cancer Genome Atlas (TCGA) cancer tissue samples, while negative samples included the adjacent normal tissues and GTEx normal tissue samples. A total of 44,673 features from the Tumor-SRT expression data were used for model training. Then we implemented five machine learning algorithms—Random Forest, Decision Tree, XGBoost ^16^, LightGBM ^17^, and CatBoost ^18^ —each optimized through grid search to determine the best hyperparameters. During grid search, 5-fold cross-validation was employed, using the Receiver Operating Characteristic Area Under the Curve (ROC AUC) as the evaluation metric. Among these models, LightGBM demonstrated the highest performance across training and test datasets, with evaluation metrics including AUC, accuracy, sensitivity, and specificity **(table S1)**. Specifically, for the test set, the model achieved an accuracy of 0.9897, sensitivity of 0.9978, specificity of 0.9665, and an AUC of 0.9995 **(Fig. 1**). To assess model robustness, we conducted validation using an independent dataset, which yielded an AUC of 0.8703 (95% CI: 0.8579-0.8813), accuracy of 0.842, sensitivity of 0.887, and specificity of 0.603, further supporting the model’s reliability **(Fig. 1**).

**Fig. 1.**
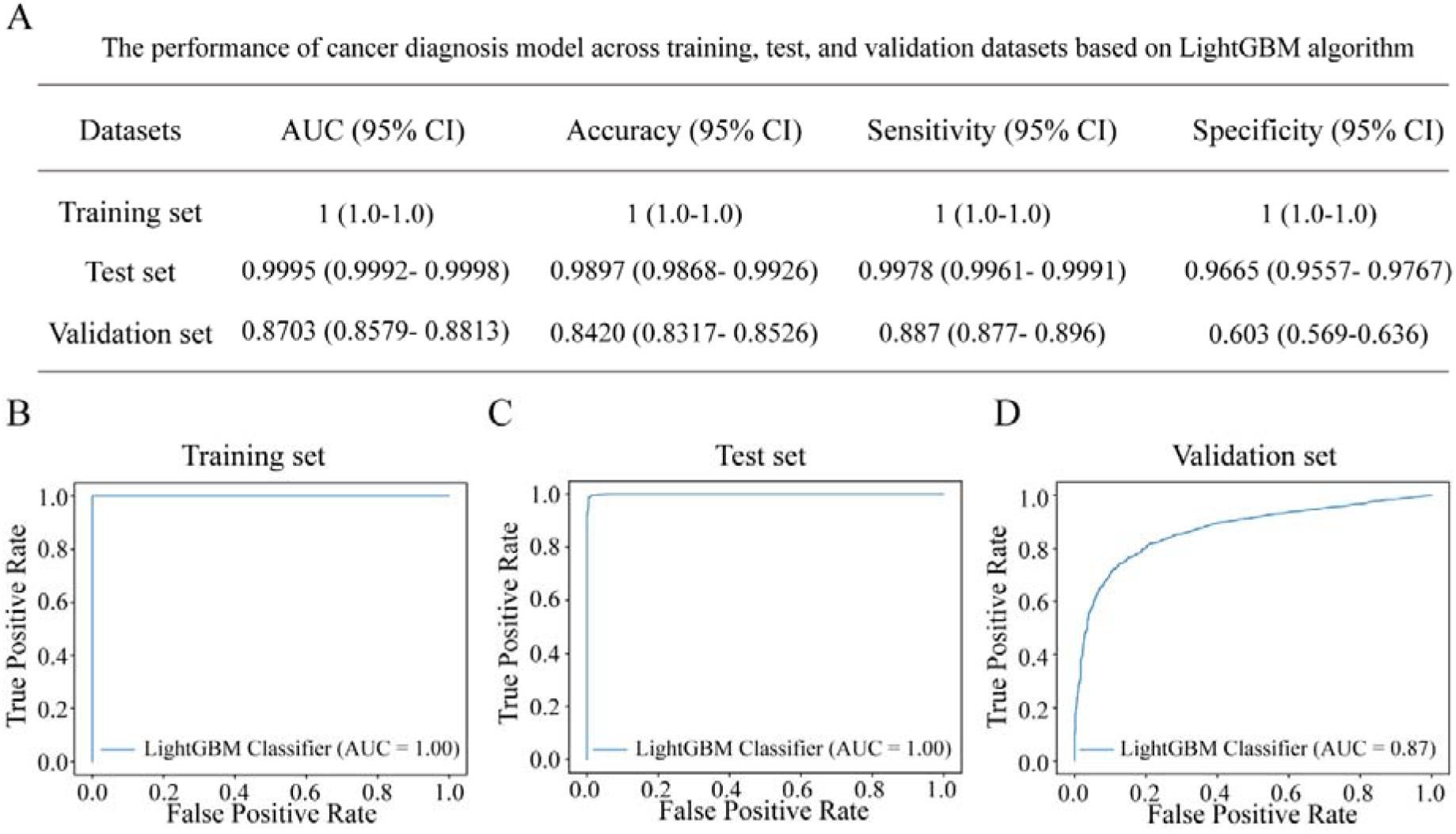
The performance of cancer diagnosis model across training, test, and validation datasets. (A) The performance of cancer diagnosis model across training, test, and validation datasets. (B-D) ROC curves illustrate the trade-off between sensitivity (true positive rate) and specificity (false positive rate) at various classification thresholds across the training (B), test (C), and validation (D) datasets. The AUC values quantify the model’s ability to distinguish between positive and negative classes, with higher values indicating better predictive performance. The LightGBM algorithm was employed to optimize the model’s accuracy and generalizability. ROC, receiver operating characteristic; AUC, area under the receiver operating characteristic curve.

### Machine Learning Models for Cancer Tissue Origin Identification

Given the tissue-specific expression of SRTs ^8,19^, we hypothesized that they could aid in tissue origin identification. We tested this by analyzing 33 TCGA cancer types, grouped into 26 categories based on their organ of origin **(table S2)**, including tissues such as the adrenal gland, B cells, bladder, brain, breast, bone & muscle, bone marrow, cervix, colon, esophagus, eye, ganglion, kidney, liver, lung, oral, ovary, pancreas, prostate, skin, stomach, testis, thymus, uterus, and pleura & peritoneum. Tissue-SRT signatures from 25 normal tissues (excluding pleura and peritoneum due to the lack of corresponding normal RNA-seq data) were used to construct machine learning classifiers for cancer tissue origin identification. Positive samples were assigned based on TCGA’s known tissue origins, while negative samples originated from other tissues. The dataset was randomly split into training and test sets at a 3:1 ratio. Five machine learning algorithms—Random Forest, Decision Tree, XGBoost, LightGBM, and CatBoost—were employed for each of the 25 tissue types (see Materials and Methods for further details). Model performance was evaluated using AUC, accuracy, sensitivity, and specificity **(table S3)**. Both training and test sets showed high predictive performance across all five algorithms, with no significant differences in performance metrics such as AUC, accuracy, sensitivity, and specificity.

To validate the model’s generalization capacity, we conducted validation using an independent dataset. Hyperparameters optimized during the training phase were applied to evaluate the 25 tissue origin models using the validation set. The model performance was quantified based on AUC, accuracy, sensitivity, and specificity, with detailed results presented in **table S3**. Notably, performance varied significantly across the 25 tissue origin identification sub-models (**Fig. 2**, P < 0.05). Decision Tree performed the worst, while the other four algorithms exhibited varying strengths. Notably, each tissue-specific model was associated with an optimal machine learning algorithm. Specifically, models with strong predictive performance often showed favorable results across multiple algorithms, while those with weaker performance demonstrated poor results across all algorithms. This variability is likely linked to the inherent separability of features between tissue types, and further investigation is needed to elucidate the underlying mechanisms **(table S4 and Fig. 2)**.

**Fig. 2.**
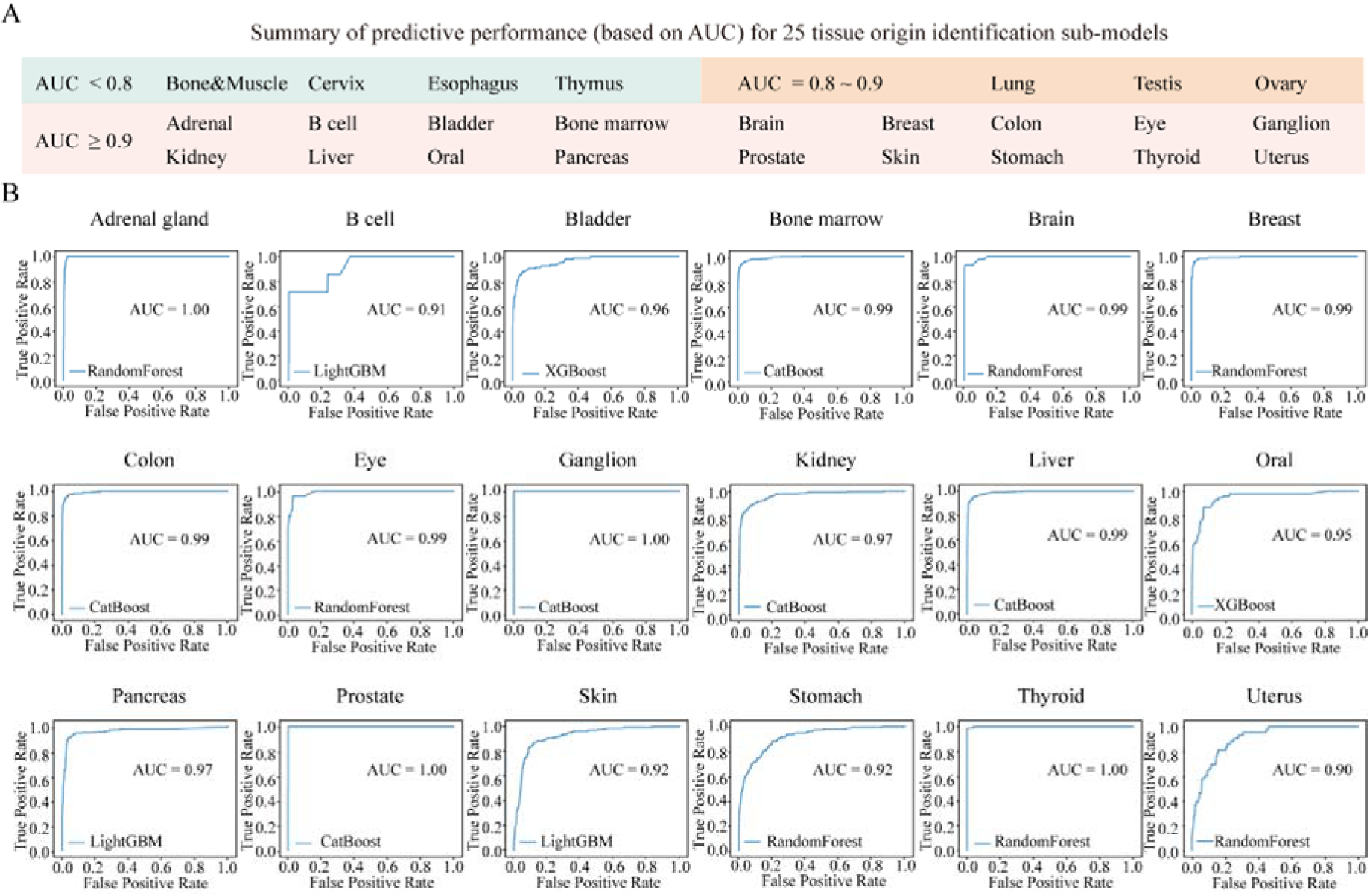
Performance evaluation of tissue origin identification models in the validation set using optimized algorithms. (A) Summary of predictive performance for 25 tissue-specific sub-models. (B) ROC curves demonstrating the sensitivity-specificity tradeoff at different classification thresholds for 18 sub-models. Each algorithm was optimized according to its corresponding tissue type. AUC values are reported for the validation cohort, demonstrating the models’ classification accuracy.

### Performance of SRT-CD in Diagnosing Cancer and Tissue Origin in Real-World Datasets

The tissue origin identification sub-models for 18 cancer types achieved an AUC greater than 0.9 with their optimal algorithms, including adrenal, bladder, bone marrow, brain, breast, colon, eye, ganglion, liver, pancreas, prostate, thyroid, B cells, kidney, head and neck, skin, stomach, and uterus (**Fig. 2A**). Building on these findings, we developed SRT-CD, an intelligent diagnostic system for precision cancer diagnosis and tissue origin identification from RNA-seq data **(Fig. 3)**. To assess its clinical applicability, we performed dual external validation using two independent datasets. First, we evaluated SRT-CD’s ability to diagnose both primary and metastatic cancers using a dataset of 209 samples: 128 hepatocellular carcinoma cases, 16 colorectal cancer cases, 15 breast cancer cases, 1 kidney cancer case, 15 gastric cancer cases, and 34 liver metastatic cancer cases (including 12 cases of breast cancer liver metastasis, 19 cases of colorectal cancer liver metastasis, and 3 cases of gastric cancer liver metastasis). When inputting the RNA-seq data of these samples into SRT-CD, the results **(Fig. 4A)** showed that the top-1 accuracy of the system in identifying the tissue origin for primary and metastatic tumors was 80% and 84.6%, respectively, meaning that in over 80% of cases, the cancer diagnosis model correctly identified the cancer’s origin on the first attempt. Furthermore, the overall top-3 accuracy was 98.1%, indicating that the system accurately identified the cancer’s origin within the top three results in nearly all cases. Second, we validated SRT-CD’s ability to determine the primary site of metastatic cancers, using a transcriptome sequencing dataset from 500 tumor patient samples (MET500), which includes 500 patient samples across 22 metastatic sites. We selected eight cancer types with well-established origins: prostate (90 cases), head and neck (22), adrenal (6), brain (5), bladder (16), pancreas (13), skin (13), and thyroid (4). The results demonstrated that for metastatic tumors with known n primary sites, the system achieved a top-1 accuracy of 75% and a top-3 accuracy of 90.4% **(Fig. 4B)**, further confirming its effectiveness in determining the primary sites of metastatic tumors. Despite variations in sample sizes, SRT-CD demonstrated stable predictive performance across both independent validation sets, underscoring its clinical potential in precision oncology.

**Fig. 3.**
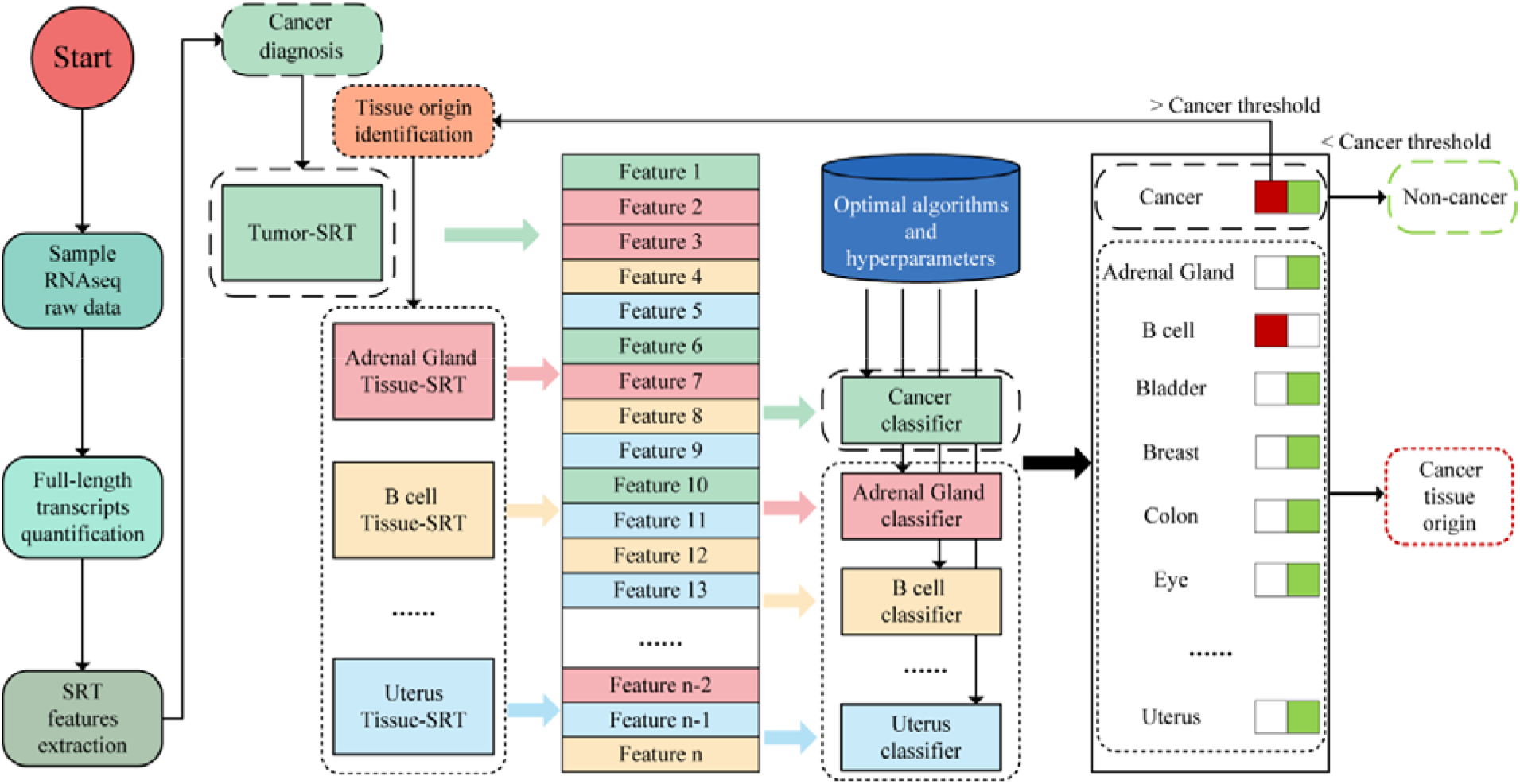
The Framework illustrates the logic procedures for diagnosing cancer and tissue origin. The framework outlines a step-by-step process for cancer diagnosis and tissue origin identification. The system consists of two main modules: a preprocessing module and a prediction module. The preprocessing module quantifies full-length RNA transcript expression from the RNA-seq data of the sample. The prediction module extracts features from the processed RNA transcript expression data and inputs them into the cancer and tissue origin identification sub-models. The SRT-CD system operates with pre-set parameters. During sample prediction, the process begins with an initial assessment by the cancer diagnostic model. Samples with prediction probabilities meeting or exceeding the predefined cancer threshold are classified as positive, while others are designated as negative. For positive samples, SRT-CD proceeds to tissue origin identification using 18 independent sub-models. Each sub-model calculates a probability value for its designated tissue origin and determines. A sample is classified as positive for a specific tissue if its probability meets the predefined threshold; otherwise, it is classified as negative. The final output includes both the raw probability values for each tissue origin and binary classification results based on the preset thresholds.

**Fig. 4.**
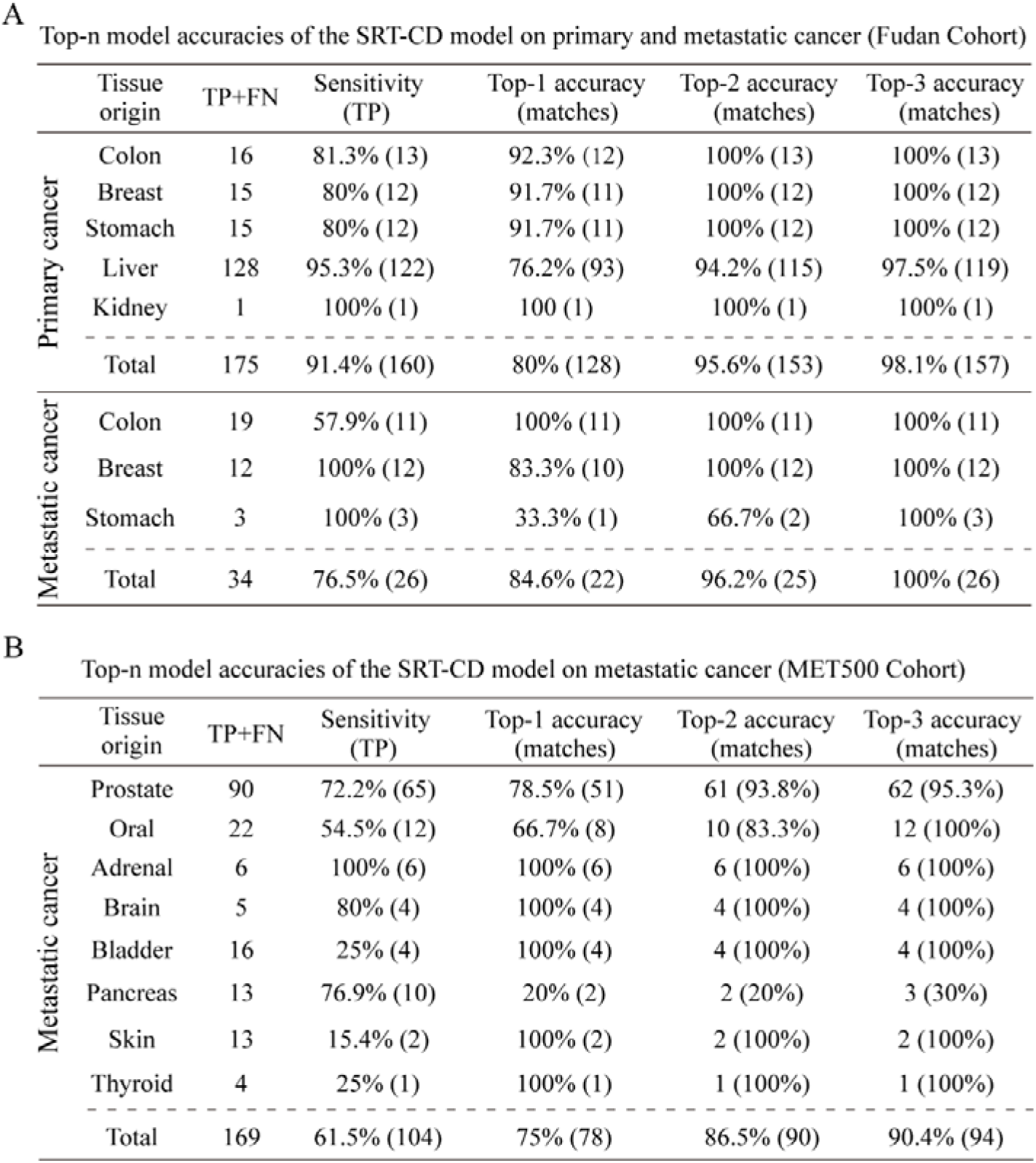
Cancer Tissue Origin identification performance of the SRT-CD model on primary and metastasis tumors. (A) Top-n model accuracies of the SRT-CD model on primary and metastasis tumors (Fudan Cohort). (B) Top-n model accuracies of the SRT-CD model on only metastasis tumors (MET500 Cohort).

## Discussion

In this study, we extracted tumor- and tissue specific RNA transcripts (SRTs) from RNA sequencing data and developed SRT-CD, a machine learning model with strong potential for distinguishing malignant tumors from benign diseases and inferring cancer provenance. Various machine learning methods have been proposed to diagnose cancer and predict tissue-of-origin using data from diverse molecular techniques, including DNA sequencing (targeted, whole exome, and whole genome), RNA profiling (coding RNA, microRNA, and whole transcriptome profiling), methylation profiling, and digital pathology ^20^. These methods indicate that machine learning approaches hold significant potential for advancing the diagnosis and treatment of CUP ^14,15,21^. However, their clinical application remains limited ^20^. In the study, we focus on a detailed analysis of SRTs across diverse human cancers and normal tissues, offering a more precise understanding of transcript expression than traditional differential gene expression analysis. By examining full-length transcripts, we identify subtle but crucial differences in expression between cancerous and normal cells, as well as among normal tissues. Moreover, this approach also enables the discovery of novel biomarkers overlooked by conventional methods, potentially leading to improved diagnostics and targeted therapies for cancer patients.

The large-scale training and testing datasets cover 33 cancer types and 22 normal tissue types from the GTEx and TCGA databases that ensure the broad applicability and generalizability of the feature extraction process. Both the cancer diagnosis model and the tissue origin identification model demonstrated high accuracy and specificity across both the training and testing sets, with AUC values exceeding 0.9 for all five machine learning models, underscoring the excellent performance of the model. Additionally, an independent external validation cohort including RNA-seq data for 33 cancer types that fully matched the training set further validated the generalizability of this system. This external validation cohort was sourced from a multicenter global dataset, comprising patients from the Americas (United States and Canada), Europe (Sweden, Poland, Germany, France, etc.), and Asia (China, South Korea, etc.), ensuring a large and diverse patient population. The validation results showed that the system achieved satisfactory predictive performance on this external dataset. A classification algorithm in machine learning is designed to identify patterns and relationships in labeled data. Two main types of classification algorithms are used: binary and multiclass. Our model uses a binary classification approach, first determining whether a sample is cancerous and then identifying its tissue origin. As third-generation sequencing data for normal tissues, cancer tissues, and adjacent tissues continue to accumulate, the amount of full-length transcript information will become more comprehensive. These models are designed to operate independently, which allows for individual optimization and adjustments, as well as flexibility for future functional expansion and performance enhancement.

Although the tumor diagnostic system demonstrated excellent performance in the training and test sets (AUC values approaching 1), its performance slightly declined in the external validation set, where the AUC value was 0.87. This performance gap is primarily attributed to the lower specificity leading to a certain risk of misclassification, where some benign lesions were incorrectly classified as malignant tumors. The limitation likely stems from an insufficient number of normal tissue samples in the training set, which may have hindered the model’s ability to fully characterize tumor-specific RNA transcripts. Future improvement should expand the range of normal tissue sample collections, particularly including more distinct tissue types. In addition, tissue origin identification sub-models varied across tumor types. While 18 tissue origin identification sub-models achieved AUC values above 0.9 in the validation set, others were prone to errors. Varying machine learning algorithms did not significantly improve the prediction performance for these tumor types, suggesting that some tissue-specific RNA transcripts were underrepresented in the training set. The model’s accuracy is heavily reliant on the representativeness of the training data and the feature composition, rather than on specific machine learning methods. More specific feature data including rare and underrepresented cancer samples will further enhance the model’s predictive capabilities. Notably, due to the absence of publicly available CUP-specific datasets (e.g., in TCGA or PCAWG), we evaluated our model on metastatic tumors. Despite the complexity of these cases, SRT-CD exhibited reliable predictive ability, with top-1 accuracy ranging from 75% to 84.6%, further surporting its clinical application potential.

In conclusion, this study highlights the capability of tumor-SRTs and tissue-SRTs as reliable biomarkers for cancer diagnosis and tissue origin identification across various cancer types. We achieved high accuracy in both tumor classification and tissue origin identification using machine learning algorithms, and developed the SRT-CD system to further support the clinical applicability. These results provide compelling evidence for the utility of SRTs in cancer diagnosis, offering a promising tool for improving cancer classification and guiding treatment decisions. We are currently sponsoring an Investigator-Initiated Trial (IIT) for diagnosing CUP using SRT-CD system and aim to implement this diagnostic system in clinical practice in the near future.

## Materials and methods

### Study Cohorts

The data used in this study were divided into training, test, validation, and external validation sets, consisting of four different cohorts.

1, The First Cohort (TCGA and GTEx Datasets, training and test sets): This cohort includes RNA-seq data from 33 types of cancer and adjacent normal tissues in the TCGA dataset ^22^, as well as normal tissue RNA-seq data from the GTEx project ^23^. These data are primarily used for model training and testing. Table S5 and Fig. S1 provides the number of patients for each of the 33 different cancer types in the American cohort (TCGA). Table S6 provides the number of samples for each of the normal tissue in the GTEx.

2, The Second Cohort (GEO and dbGAP Datasets, validation sets ^24,25^): This cohort includes RNA-seq data from 33 types of cancer and adjacent normal tissues, obtained from public databases, including Gene-Expression Omnibus (GEO) and dbGAP. These data are primarily used for model validation. Table S7 and Fig. S1 provides the number of patients for each of the 33 different cancer types in the American cohort.

3, The Third Cohort (External Evaluation Set): This external cohort is used to assess the accuracy of the developed diagnostic model for distinguishing between primary and metastatic cancers. It includes data from 128 cases of hepatocellular carcinoma, 16 cases of colorectal cancer, 15 cases of breast cancer, 1 case of renal cancer, 15 case of gastric cancer, and 34 cases of liver metastatic carcinoma (Fig. S2). These datasets were sourced from GepLiver (www.GepLiver.org/#/download) ^26^ and HRA003557 (https://ngdc.cncb.ac.cn/gsa-human/) ^27^.

4, The Fourth Cohort (External Evaluation Set from MET500 ^28^): This cohort is used to evaluate the accuracy of the developed model in predicting the primary site of metastatic cancers. It consists of transcriptome sequencing data from 500 tumor patients with a total of 22 organ-specific metastatic lesions. Based on the accuracy of the cancer diagnostic model, 13 known cancer types (adrenal, breast, colon, liver, brain, prostate, thyroid, bladder, head and neck, kidney, pancreas, skin, and stomach) were selected for prediction (Fig. S2). This study utilized publicly available de-identified datasets and was exempt from institutional review board approval.

### RNA-seq Data Collection and Processing

RNA-seq data across 33 different human cancer types were retrieved from TCGA, GEO, and dbGAP. RNA-seq data from 25 different human normal tissue types were downloaded from GTEx and GEO. BAM files were converted to FASTQ format using Samtools (version 1.7). Transcript expression, measured in transcripts per kilobase million (TPM), was quantified by mapping RNA-seq reads to long-read isoforms using Salmon (version 1.5.2). The RNA-seq data from training, testing, and validation datasets were used to quantify the expression of full-length RNA transcripts, thereby obtaining specific RNA transcript expression levels for the respective tissue samples.

### Building the Diagnostic SRT Signature

Tumor-specific RNA transcripts (Tumor-SRTs) detected in LR-seq are defined as those found exclusively in tumor samples and not in normal tissues. The criteria for identifying Tumor-SRTs through RNA-seq include: 1, The median expression level of the transcript in tumors is at least 10 times higher than the maximum expression level observed in all normal tissues (excluding testis) in the GTEx dataset, as well as adjacent normal tissues in the TCGA dataset; 2, The transcript is expressed (TPM > 0.5) in >5% of tumor samples from at least one TCGA cancer type; 3, The methodology for defining tissue-specific RNA transcripts (tissue-SRTs) follows a similar approach to that described in a previous study. Tumor-SRT and tissue-SRT data were downloaded from the FLIBase database^8^.

### Training and validation for Cancer Diagnosis

Cancer tissue samples from the TCGA dataset served as positive samples, while adjacent normal tissue samples and all normal tissue samples from the GTEx dataset were used as negative samples. The positive and negative samples were randomly partitioned into training and testing sets in a 3:1 ratio to ensure the independence of the two datasets.

Training was conducted using the training set data and various machine learning models, including Random Forest, Decision Tree, XGBoost, LightGBM, and CatBoost. The training features were based on the quantitative results of 44,673 tumor-specific RNA transcripts, represented by TPM values. Each sample in the training set included the TPM values of these 44,673 tumor-specific RNA transcripts, which were input into the machine learning models for model development. Hyperparameter optimization was performed using grid search to determine the optimal settings for each model. During the grid search, each parameter combination was evaluated using 5-fold cross-validation, with the ROC AUC serving as the performance metric. The AUC value of 0.9-1.0 is excellent, 0.8-0.9 is good, 0.7-0.8 is average, 0.6-0.7 is poor, and 0.5-0.6 is very poor. This process yielded the hyperparameters that optimized model performance. Five cancer diagnosis models were generated, corresponding to the aforementioned machine learning algorithms.

The performance of the five cancer diagnosis models was validated on an independent validation set, using evaluation metrics that included AUC, accuracy, sensitivity, and specificity. The model exhibiting the highest performance across these metrics was selected as the final cancer diagnosis model. The validation set consisted of RNA-seq data from 33 cancer types and their corresponding adjacent normal tissue samples, obtained from public repositories such as GEO and dbGaP. In this dataset, cancer tissue samples from all GEO and dbGaP datasets were treated as positive samples, while adjacent normal tissue samples were classified as negative samples.

### Training and validation for Cancer Tissue Origin Identification

Data for training and testing were sourced from 33 types of cancer tissues in the TCGA dataset. These cancer tissue samples were categorized according to both tissue origin and cancer type, which included the following tissue types: adrenal gland, B cells, bladder, brain, breast, bone & muscle, bone marrow, cervix, colon, esophagus, eye, ganglia, kidney, liver, lung, head and neck, ovary, pancreas, prostate, skin, stomach, testis, thymus, thyroid, and uterus, making a total of 26 tissue categories.

The features for model training consisted of the quantitative expression values TPM of 25 distinct normal tissue-specific RNA transcripts. The number of RNA transcripts for each tissue is outlined in table S2. These features were employed to construct corresponding sub-models for cancer tissue classification. Due to the absence of RNA-seq data for peritoneal tissue, it was excluded from model predictions. A sub-model was constructed for each of the 25 tissues to predict whether a test sample originates from that specific tissue type. In the training process, cancer samples with a known tissue origin (from TCGA) were assigned as positive samples, while those from other tissue origins were classified as negative samples. Training and testing sets were partitioned in a 3:1 ratio, and a binary classification approach was used. This process was repeated for each tissue type, yielding 25 sub-models, each designed to classify whether a sample originates from a specific tissue type.

For example, on the text of the adrenal tissue origin identification sub-model, cancer samples originating from the adrenal gland (e.g., adrenal cortical carcinoma) were considered positive, while samples from other tissue origins were treated as negative. A 3:1 split was used for training and testing, and hyperparameter optimization was conducted using grid search with 5-fold cross-validation. The ROC AUC served as the scoring function for selecting the optimal hyperparameters. The machine learning algorithms used for this task included Random Forest, Decision Tree, XGBoost, LightGBM, and CatBoost. The process for constructing the remaining 24 tissue origin identification sub-models followed the same methodology outlined for the adrenal model. In total, 25 tissue origin identification sub-models were developed, each comprising five machine learning model predictions. These sub-models were used to determine whether a test sample originated from any of the 25 tissues.

The performance of the 25 tissue origin identification sub-models was validated using an independent validation dataset. Evaluation metrics included AUC, accuracy, sensitivity, and specificity. The sub-model with the best overall performance was selected as the final tissue origin identification sub-model. The validation dataset was derived from RNA-seq data of 33 cancer tissue types, obtained from public repositories such as GEO and dbGaP. Based on the results obtained from the 25 tissue origin identification sub-models, the tissue origin identification models for 18 types of cancer achieved an AUC greater than 0.9 for their optimal algorithms. These cancers include adrenal, bladder, bone marrow, brain, breast, colon, eye, ganglion, liver, pancreas, prostate, thyroid, B cells, kidney, head and neck, skin, stomach, and uterus.

### Developing Specific RNA Transcript-based Cancer Diagnostics (SRT-CD)

Based on this performance, a software tool was developed to diagnose cancer using the specific transcript data expressed by the cancer patient’s tissue. Additionally, the software determines whether the cancer originates from one of the 18 tissues. The system comprises two modules: a preprocessing module and a prediction module. The preprocessing module is responsible for quantifying the full-length RNA transcript expression based on RNA-seq raw data of the sample to be tested. The prediction module extracts the features from the full-length RNA transcript expression data and inputs them into the cancer and tissue source identification models. The parameters for the identification models are pre-set in the system, allowing the software to predict whether the input sample is cancerous and to determine the probability and threshold for each tissue origin. We used top-n accuracy to evaluate the performance of tissue origin identification, and set n as 1, 2 and 3. Top-1, -2 and -3 accuracy was used to measure frequency in regard to the correct label found, and to make the maximum confidence prediction. Top-n accuracy looks at the nth classes with the highest predicted probabilities when calculating accuracy. If one of the top-n classes matches the ground-truth label, the prediction is considered to be accurate.

### Statistics

Bootstrapping, implemented in Python, was used to compute confidence intervals for accuracy, sensitivity, specificity, and ROC AUC. These four metrics were calculated using the accuracy_score, recall_score, and roc_auc_score functions from the sklearn package (v.1.3.1). For multi-class ROC AUC, the one-vs-rest strategy was used for calculation. Machine learning algorithms, including Random Forest, Decision Tree, XGBoost, LightGBM, and CatBoost, were trained on the training set and tuned via grid search for optimal hyperparameters. Hyperparameter optimization was performed using 5-fold cross-validation, with the ROC-AUC serving as the primary evaluation metric.

For external validation, independent datasets were used to assess the generalizability of the models. The models were applied to the external datasets using the optimized hyperparameters, and their predictive accuracy was evaluated based on AUC, accuracy, sensitivity, and specificity. The performance of the models was statistically compared using paired t test to assess significant differences in performance metrics (P < 0.05). For evaluation of the SRT-CTC system, we computed top-1 and top-3 accuracy metrics, where top-1 accuracy reflects the proportion of samples for which the model’s top prediction was correct, and top-3 accuracy refers to the proportion of samples where the true label was included within the top three predictions. Statistical significance for all comparisons was set at P < 0.05. All results are presented with their corresponding 95% confidence intervals where applicable. Python version 3.10.12 was used for all analyses.

## Code availability

We used Python (v3.10.12) for classification and analysis, utilizing machine learning algorithms from the scikit-learn package ^29^ (v.1.3.1), CatBoost ^18^(v.1.2.3), XGBoost ^16^(v.2.0.3), and LightGBM ^17^(v.4.3.0). These libraries were employed for model development and classification tasks.

## Author contributions

Conceptualization: X.H., Z.C., Y.C. Methodology: Z.C., J.T., H.L.

Visualization: X.H., Z.C., Y.C., H.L. Funding acquisition: X.H., Z.C. Project administration: X.H., Z.C. Supervision: X.H., Z.C., Y.C. Writing – original draft: Z.C., J.T.

Writing – review & editing: X.H., Y.C.

All authors read and approved the final manuscript.

